# *Tamarix articulata* extract offers protection against toxicity induced by beauty products in Hs27 human skin fibroblasts

**DOI:** 10.1101/2023.05.30.542982

**Authors:** Abdullah M Alnuqaydan, Faten M. Ali Zainy, Abdulmajeed G Almutary, Najwane Said-Sadier, Bilal Rah

## Abstract

The current study evaluates the cytotoxicity and chemical analysis of selected beauty products and evaluation of the protective effect of *Tamarix articulata* (TA) extract against toxicity induced by beauty products in skin fibroblasts (Hs27). MTT and Crystal violet (CV) assays were used to determine the dose-dependent cytotoxic effects of products against Hs27 fibroblasts. DNA fragmentation assay was conducted to determine the mode of cell killing induced by evaluated beauty products. Chemical analysis and heavy metals were evaluated to determine beauty products. Pre-treatment with TA extract for different time points followed by time-dependent exposure with beauty products to assess the protective effect of TA extract in Hs27 cells was analysed by MTT and CV assays. Owing to the presence of various harmful heavy metals such as arsenic (As), chromium (Cr), cadmium (Cd), nickel (Ni), and lead (Pb) in beauty products, our results revealed that all beauty products induce significant cytotoxicity over time (1, 4 h) in a dose-dependent (125, 250, 500 µg/mL) manner. DNA fragmentation assay revealed that the induced cytotoxicity was caused by necrosis. However, pre-incubation with a safe dose (50 µg/mL) of TA for different times (24, 48 h) followed by exposure to various doses (62.5, 125, 250, 500 µg/mL) of beauty products for different times (1, 4 h) revealed significant (\**p*≤0.05, \*\**p*≤0.01) protection against product-mediated cytotoxicity. The effect was more pronounced for 1 h exposure to products compared to 4 h. Our study demonstrates that the due to the presence of heavy metals in synthetic beauty products exhibits marked toxicity to skin fibroblasts. However, the presence of abundant bioactive polyphenols with promising antiscavenging activity in TA extracts significantly nullifies cytotoxicity promoted by examined beauty products in skin fibroblasts (Hs27).

## Introduction

Skin is the largest organ and covers externally the whole human body to regulate temperature while fluid balance restricts the dangers posed by agents including chemicals, microbes, and protects against harmful sunlight-driven ultra-violet (UV) radiation. Long-term exposure to radiation (ionizing and UV), drugs, chemicals, and cosmetic products including personal care products containing chemicals will harm the human skin and other organs of the body after absorption [1]. Continuous exposure to these products and radiation has serious implications such as lethal and prolonged absorption and life-threatening anaphylaxis (allergies) [2]. Apart from these outcomes, they can generate reactive oxygen species (ROS) in cutaneous cells and subsequently triggers the induction and activation of oxidative stress pathways [3]. This uncontrolled generation of ROS has been implicated in the pathogenesis of various skin diseases and often culminates in melanoma or skin cancer [4].

Synthetic cosmetic products are the fastest-growing market around the world with claims that their active ingredients improve radiance, texture, skin tone and reduced wrinkles [5]. However, recent United States Food and Drug Administration (FDA) report suggests that more than 12,000 synthetic and related chemicals have been used in cosmetic products of which only less than 20% have proved to be non-toxic [6]. The synthetic cosmetic products after absorption into the human body through various routes can have harmful effects such as disruption of endocrine and reproductive systems, functions as carcinogens and/or neurotoxicants. In effect, they pose a serious long-term threat to human beings [7].

Before the emergence of synthetic cosmetic products, natural ingredients derived from plant sources were for centuries used as personal care products in many societies. Owing to the presence of abundant polyphenolic bioactive compounds that exhibit promising anti-scavenging activity with less toxicity and provides nutrients to skin cells [8], natural ingredients in plant-derived beauty care products provide exogenous support of antioxidants and vitamins to skin cells. They neutralize ROS and other oxidative stress-related products to keep the skin cells healthy [9]. One such plant extract ingredients derived from the family *Tamaricaceae* is *Tamarix articulata* (TA). Phytochemical analysis revealed that the TA extract is rich in polyphenolic compounds which displays promising pharmacological activities. TA is a halophytic plant abundantly found in the deserts of Saudi Arabia. Traditionally, TA has been used as a folk medicine by a tribal population – the Tafilalet in Morocco -against various conditions including skin diseases [10]. Owing to the presence of abundant polyphenolic compounds, TA extract showed promising antioxidant potential to scavenge ROS species generated during oxidative stress [11]. The current study was designed to evaluate the protection offered by TA extract against the oxidative stress-mediated toxicity induced by beauty products in human skin fibroblasts. Together, these results suggest that the presence of abundant bioactive polyphenols with promising anti-scavenging activity of TA extracts significantly nullifies cytotoxicity promoted by beauty products in skin fibroblasts.

## Materials and Methods

### Plant extract identification and characterization

*Tamarix articulata* (TA) plants parts (stem, leaves, and roots) were used in this study and were collected in December 2019 from the Qassim region of Saudi Arabia along with dried leaves found on the ground [11]. The identification and phytochemical analysis we recently published showed that major constituents of TA extract exhibits various pharmacological activities [10–13]. Further, phytochemical analysis suggests that TA extract is abundant in polyphenols **(Table 1)**. The phytochemical analysis of the methanolic extract of TA by LC-MS analysis revealed that more than 200 compounds were identified (Supplement Table 1). The key phytochemicals identified from the methanolic extract of TA by LC-MS display anticancer activities against various cellular models and are summarized in Supplement Table 1 and supplementary Figure 1.

**Table 1.**
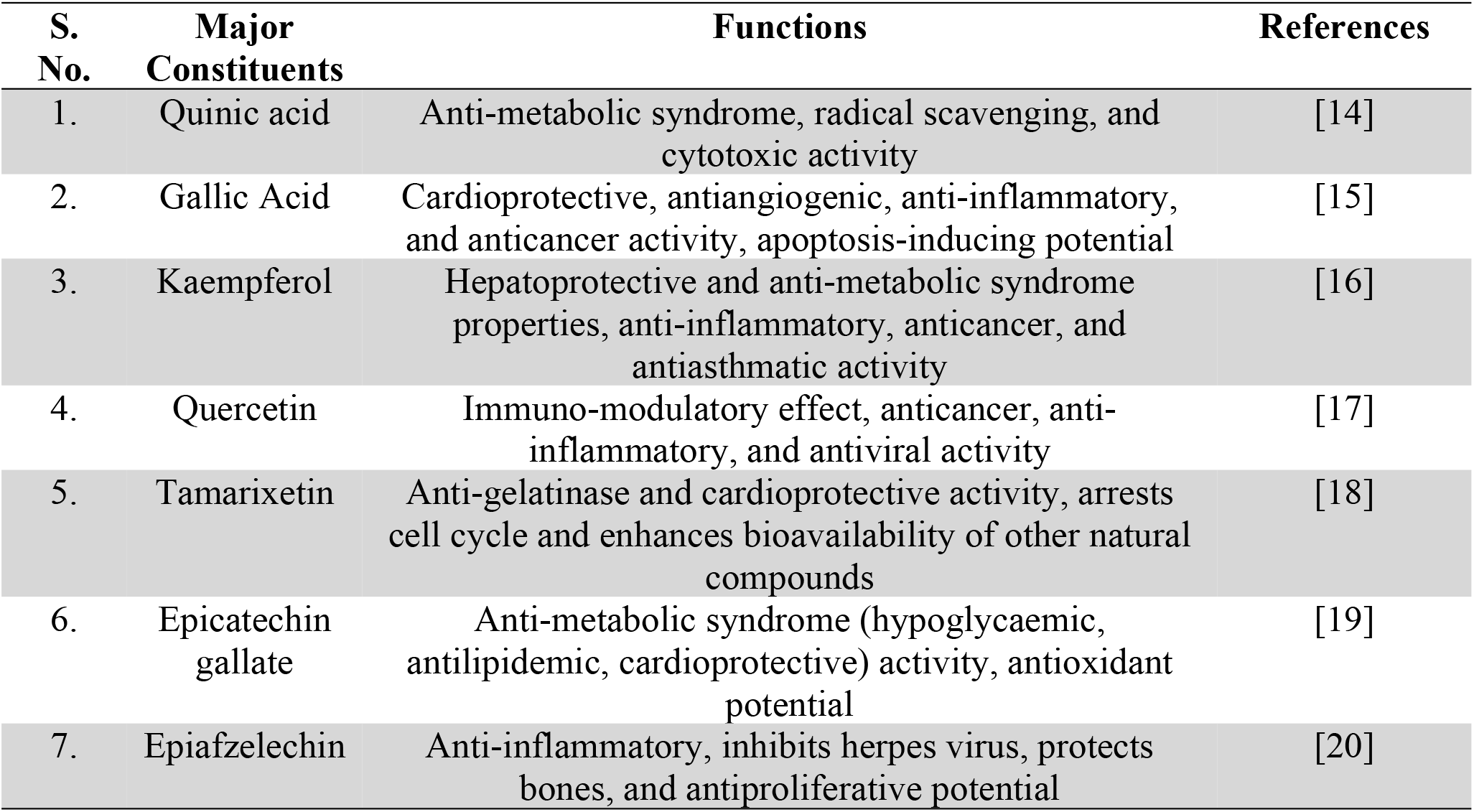
Major constituents of *Tamarix articulata* and their functions

| S. No. | Major Constituents | Functions | References |
| --- | --- | --- | --- |
| 1. | Quinic acid | Anti-metabolic syndrome, radical scavenging, and cytotoxic activity | [14] |
| 2. | Gallic Acid | Cardioprotective, antiangiogenic, anti-inflammatory, and anticancer activity, apoptosis-inducing potential | [15] |
| 3. | Kaempferol | Hepatoprotective and anti-metabolic syndrome properties, anti-inflammatory, anticancer, and antiasthmatic activity | [16] |
| 4. | Quercetin | Immuno-modulatory effect, anticancer, anti-inflammatory, and antiviral activity | [17] |
| 5. | Tamarixetin | Anti-gelatinase and cardioprotective activity, arrests cell cycle and enhances bioavailability of other natural compounds | [18] |
| 6. | Epicatechin gallate | Anti-metabolic syndrome (hypoglycaemic, antilipidemic, cardioprotective) activity, antioxidant potential | [19] |
| 7. | Epiafzelechin | Anti-inflammatory, inhibits herpes virus, protects bones, and antiproliferative potential | [20] |

### Liquid chromatography-mass spectrometry (LC–MS) metabolomic analysis and data processing

Liquid chromatography-mass spectrometry (LC–MS) was carried out as described in our recently published studies [11,21,22]. The LC–MS metabolomic analysis consisted of an AC-QUITY UPLC I-Class System (Waters Technologies, USA) coupled with a 6500 Qtrap (AB Sciex, Canada). A Zorbax XDB C18 column (2.1×150 mm. 3.5 µm) (Agilent, USA) was used for the chromatographic separation and kept at 40 °C with a flow rate of 300 µL/min. The mobile phase consisted of A (0.1% formic acid in HPLC grade water) and B (0.1% formic acid in HPLC grade acetonitrile). The linear gradient elution was as follows: 2% B (from 0 to 2), 95% B (from 2 to 24), 95% B (held for 2 min), and then 4 min equilibration time. Electrospray ionization mass spectra (ESI-MS) were acquired in the positive (ES +), with an electrode voltage of 5500 V. The delustering potential was set at 90 V and the entrance potential was 10 V. Nitrogen was used as curtain gas (30 psi) and nebulizer gas on the MS. Spectra were collected with a mass range of 100 – 900 m/z. LC data files were converted to MZxml format using MS Convert (Pro-teoWizard 3.0.20270). MZ mine software (version 2.53) was used for the analysis of the data. After importing the data into the MZ mine, a minimum intensity cut-off of 1,000 was applied and the retention time was adjusted with a tolerance of 0.2 min. Adjusted peaks were then aligned into one mass list to facilitate identification and comparison. Compounds of interest were identified by using the KEGG Database in the finalized list based on m/z with a tolerance of 30 ppm.

### Beauty products sample collection

A total of six lipstick products - three products from famous brands Huda Beauty, Revlon, and Maybelline (Saudi market – August-2019) and other three are unbranded - were collected from a local market. Beesline lipstick product served as the standard sample (made in Lebanon). The analysed original brand lipstick samples were manufactured in Italy, USA, and Paris, while the unbranded lipstick products taken from the market have unknown manufacturer source. The prices of the original brand lipstick samples ranged from $28.00 to $11.37 USD per sample, while the unbranded samples ranged from $3.00 to $2.00 USD. All the samples were transferred to the laboratory for the analysis of their cytotoxicity and heavy metals.

### Chemicals and reagents

The cell culture media Rosewell Park Memorial Institute (RPMI)-1640, antibiotics (penicillin-streptomycin), phosphate-buffered saline (PBS), and fetal bovine serum (FBS) were procured from Invitrogen. DNA fragmentation assay kit and MTT were procured from Abcam. All other reagents and chemicals required for assays were purchased from Sigma-Aldrich.

### Cell culture

Human skin fibroblast cell line Hs27 cells were procured from ATCC. The cell line was maintained in culture media (Rosewell Park Memorial Institute (RPMI)-1640) supplemented with 10% FBS and 1% penicillin-streptomycin to avoid any bacterial contamination. The cells were grown and incubated in a 5% humidified CO_2_ incubator in sterile tissue culture T75 flasks. Hs27 cells were periodically checked for Mycoplasma contamination.

### Cell treatments

Human skin fibroblasts Hs27 cells were cultured overnight to adhere to the surface of a flat bottom 96-well microplate at a density of 10_4_ cells per well and incubated in a 5% humidified CO_2_ incubator at 37 °C. Next morning culture media was aspirated from each well of the microplate and replaced with fresh media containing varying doses (62.5, 125, 250, and 500 µg/mL) of personal care products for 1, and 4 h to determine cytotoxicity. However, for protection assay Hs27 cell after overnight incubation, cells adhered in wells of the 96-well microplate were preincubated with non-toxic doses (50 µg/mL) of TA extract for different times (24, 48 h). This was followed by exposure to varying doses (62.5, 125, 250, and 500 µg/mL) of personal care products for 1 h and 4 h to determine the level of protection against personal care product-induced cytotoxicity.

### Cell viability (MTT) assay

The cell viability of human skin fibroblasts (Hs27) was determined by MTT assay as per the standard protocol [23]. Briefly, 10_4_ Hs27 cells were seeded per well in the 96-microplate well which allowed the cells to adhere to the microplate’s bottom surface. The culture media in each well was aspirated and had fresh media added to it, containing varying doses of personal care products (62.5, 125, 250, and 500 µg/mL) and a non-toxic dose of TA extract (50 µg/mL) for different times (1, 4 h) and (24, 48 h), respectively. Following the completion of times, the wells containing cells incubated with 20 µl MTT dye (2.5 mg/mL) for 3-4 h at 37 °C in 5% CO_2_ incubator. The formazan crystals formed during incubation were dissolved in DMSO (150 µl) by gentle vortex to ensure complete dissolution. The purple-colored solution formed in each well was measured at 570 nm by multiplate reader.

### Crystal violet (CV) staining for cell viability

Briefly, 10_4_ cells were seeded in each well of the 96-microplate well to determine cell viability utilizing CV assay. After overnight incubation to the cells and properly adhering to the bottom surface of the plate well, Hs27 skin fibroblasts were exposed to different concentrations of beauty (lipstick) products for varying times (1 and 4 h). The cells in each well were washed with PBS gently and then incubated with 50 µL of 0.5% CV solution in methanol for 10 min at room temperature. The wells of the 96-microplate well plate was washed with distilled water and then air-dried. After the plate wells were air-dried 50 µL distaining solution (33% acetic acid) was added. The blue color solution thus obtained was measured by multiplate reader at a wavelength of 570 nm.

### Sample preparation and analysis

The sample preparation for heavy metal analysis was done according to the standard method [24,25] by using ICP-OES (Thermo-Scientific; ICAP 6000 Series). The standard operational conditions for the ICP-OES work were as follows: 1550 W-power; 15 L/min-plasma gas; 0.2 L/min-aux gas; 0.8 L/min-nebulizer; and 0.3 mL/min-sampling rate [26,27]. To compare the quality control of original and fake brands HNO_3_ and HClO_4_ (65% - 60%, Sigma-Aldrich) were adjusted so that they could digest the lipstick samples [28]. All the solutions were prepared in double-distilled water and required dilutions were created for analysis. Five heavy metals were investigated: As, Cd, Cr, Pb, and Ni. Every day, fresh calibration standards for each metal were prepared from the certified standard stock solution (High Purity Standards ICP-OES-68B Solution A, 100 mg/L in 4% HNO3) in the 0.5 to 10 ppm range.

### Apoptosis DNA ladder assay

The assay served to determine whether the cytotoxicity induced by personal care products was due to apoptosis or necrosis. Briefly, 0.5 ×10_6_ Hs27 cells were exposed to certain doses (250 and 500 µg/mL) of personal care products for 4 h and then harvested. DNA isolation of treated cells was done as per the manufacturer’s instructions stated in Apoptosis DNA Ladder Assay kit purchased from Abcam. After the isolation of genomic DNA from cells exposed to varying doses of personal care products, the samples from each treatment were resolved and analysed by 2% agarose gel electrophoresis containing ethidium bromide (EtBr). After resolving was completed the agarose gel was analysed by gel doc to generate an image.

### Statistical Analysis

All the experiments reported in the current study were done in triplicate. The results of the current study were calculated, processed by one-way ANOVA, and represents the mean of ± SEM. The *p*-value equal to or less than 0.05 was deemed to be significant (* means *p* ≤ 0.05, ** means *p* ≤ 0.01 and *** mean *p* ≤ 0.001).

## Results

### Liquid chromatography-mass spectrometry (LC–MS) metabolomic analysis and data processing

Liquid chromatography-mass spectrometry (LC–MS) was carried out as described in our recently published studies [11,21,22]. The finding of phytochemical analysis of the methanolic extract of TA performed by LC-MS analysis revealed that more than 200 compounds were identified (Supplement Table 1). The key phytochemicals identified from the methanolic extract of TA by LC-MS display anticancer activities against various cellular models and are summarized in Supplement Table 1 and supplementary Figure 1.

### Cytotoxic effects of personal care products on human skin fibroblasts (Hs27)

The cytotoxic effects of personal care products were determined by the MTT assay against the *in-vitro* skin cellular model (Hs27 cells). The MTT assay evaluated the cell viability against varying doses of personal care products. After seeding, Hs27 cells at a density of 104 cells per well were put into 96-well microplates and adhered properly to the bottom surface overnight at 37 °C in a 5% humidified CO2 incubator. Hs27 cells in the 96 wells were exposed to varying doses (62.5, 125, 250, and 500 µg/mL dissolved in culture media) of personal care products for lengths of time (1, 4 h) along with untreated control and a standard compound, which is known to exert an insignificant cytotoxic effect on human skin fibroblasts due to the absence of heavy metals and other toxic chemicals.

Our MTT assay results demonstrate that significant cytotoxicity (*p* ≤ 0.05) was induced by personal care products at all doses of personal care products and times (1, 4 h) treatment when compared with untreated control and a standard compound. The latter revealed less cytotoxicity (Fig. 1a, b). To verify the above MTT assay results, we undertook the crystal violet (CV) assay. Intriguingly, we observed a similar pattern with a significant percentage of cell cytotoxicity of Hs27 cells at all doses (62.5, 125, 250, and 500 µg/mL) of personal care products and times (1, 4 h) as found earlier with the MTT assay However, we did not observe any significant cytotoxicity of Hs27 cells when exposed to different doses of a standard compound at all times when compared to untreated control (Fig. 1a, b). Together, the cytotoxicity results suggest that: firstly, personal care products induce significant cytotoxicity to human skin fibroblasts Hs27; and secondly, might have induced and activated oxidative stress pathways to kill cells.

**Fig. 1.**
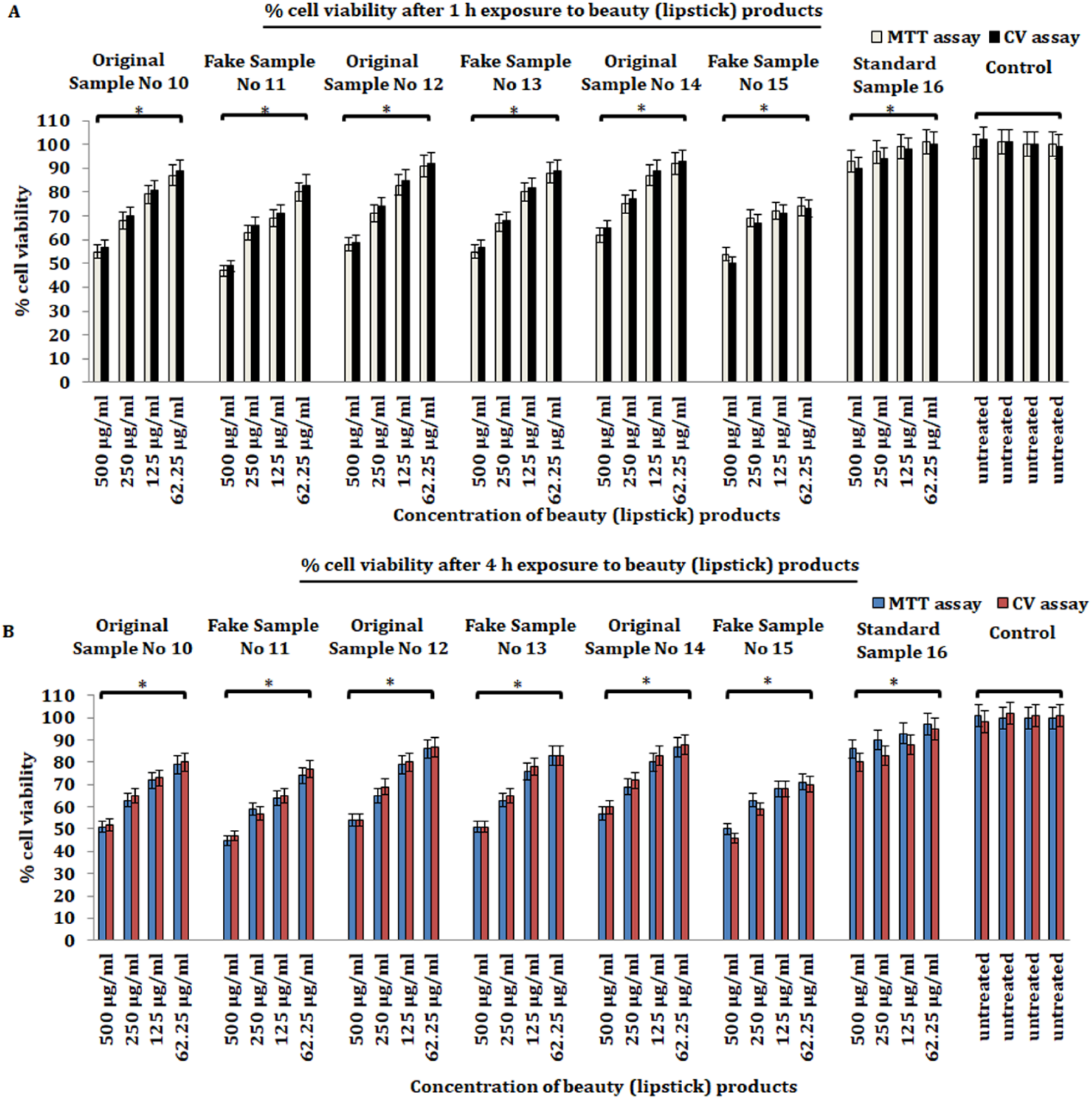
Evaluation of cell viability of Hs27 skin fibroblasts. (**a**) Percent of cell viability of Hs27 skin fibroblasts exposed to varying concentrations (62.5, 125, 250, 500 µg/mL) of personal care products for 1 h (**b**) Percent cell viability of Hs27 skin fibroblasts exposed to varying concentrations (62.5, 125, 250, 500 µg/mL) of personal care products for 4 h. The data presented here is based on experiments done in triplicate and the mean value of ± SE. The p-value less or equal to 0.05 was statistically significant, \**p* ≤ 0.05.

Next, we sought to examine the mode of cell death induced by personal care products on skin fibroblasts Hs27. Following this, Hs27 cells were exposed to higher doses (250 and 500 µg/mL) of personal care products for 4 h along with untreated control and positive control camptothecin (5 µM) to analyse the mode of cell death in Hs27 cells. We executed DNA fragmentation assay by gel electrophoresis which is one of the most reliable methods for detecting apoptosis. Our results reveal that after resolving genomic DNA by electrophoresis in 1% agarose gel a smear pattern appeared, and no DNA ladder formation appeared in cell extract samples exposed to larger doses (250, 500 µg/mL) of personal care products. Nonetheless we observed a clear DNA ladder formation pattern in <u>a</u> lane of cell extract exposed to positive control camptothecin (10 µM) (Fig. 2). These findings indicated that personal care product-induced cytotoxicity was not due to apoptosis but possibly caused by oxidative stress-mediated necrosis of skin fibroblasts.

**Fig. 2.**
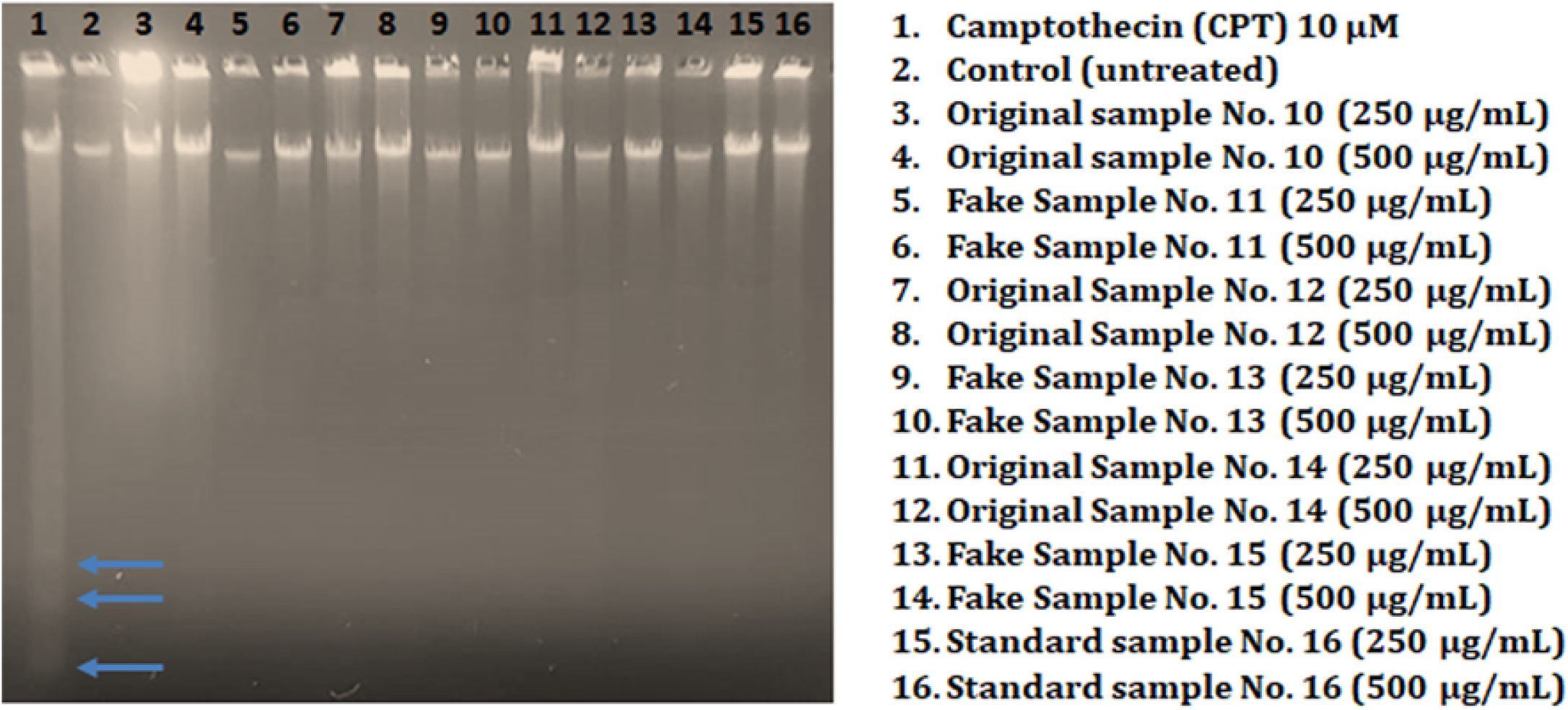
Determination of DNA fragmentation by agarose gel electrophoresis. Hs27 skin fibroblasts were exposed to varying doses of beauty (lipstick) products along with 10 µM camptothecin (positive control) to investigate the mode of cell death induced in Hs 27 cells. Arrows in the positive control lane showed DNA fragments indicate the apoptosis cell death, while the absence of DNA fragments in cells exposed to various doses of personal care products indicates that cell cytotoxicity induced by personal care products is due to necrosis.

### Chemical and heavy metal analysis of the selected beauty products

Determining the various chemicals and heavy metals in selected beauty products collected from the Saudi market as discussed in the material methods section was conducted via the ICP-OES analysis system. Five heavy metals - arsenic (As), cadmium (Cd), chromium (Cr), lead (Pb), and nickel (Ni) - were performed using concentrated perchloric and nitric acid (1:4 ratio) to analyse these heavy metals. After analysing the personal care products by the ICP-OES system, our results revealed that a small concentration of all five heavy metals was not only present in the fake beauty products (lipstick) but also in the branded ones (**Table 2**), yet the concentration was within the permissible limits as stipulated by the FDA. Interestingly, we did not find any heavy metal in the standard sample, meaning that the synthetic cosmetic products whether fake or branded (original) contain heavy metals that might induce collective toxicity to skin cells. This is well supported by significant cell killing of Hs27 skin fibroblasts by MTT assay as discussed in the previous results section.

**Table 2.** Concentrations (ppb) of heavy metals elements in beauty (lipstick) products

| Sample No. | Arsenic (As) | Cadmium Cd) | Chromium<br>(Cr) | Lead (Pb) | Nickel (Ni) |
| --- | --- | --- | --- | --- | --- |
| **Original 10** | ND* | 2 | 15 | 22 | 27 |
| **Fake 11** | ND | 2 | 15 | 22 | 27 |
| **Original 12** | ND | 9 | 25 | ND | 26 |
| **Fake 13** | ND | ND | ND | 4 | 14 |
| **Original 14** | ND | 6 | 21 | 5 | 28 |
| **Fake 15** | ND | 1 | 19 | 5 | 23 |
| **Standard**<br>**16** | ND | ND | ND | ND | ND |

\* ND: not detected

### TA extract protects against beauty products-mediated toxicity in skin fibroblasts

Owing to the presence of various polyphenols and flavonoid compounds that possess promising antioxidant properties, numerous plant products when combined with synthetic beauty products prevent the formation of ROS and other adverse effects on skin fibroblasts. We intended to determine whether TA extract could offer protection against personal care products-induced cytotoxicity to skin fibroblasts Hs27. To do this, we preincubated skin fibroblasts Hs27 with a safe dose (non-toxic) dose of TA extract (50 µg/mL) for different lengths of time (24, 48 h), followed by exposure to varying doses (62.5, 125, 250, and 500 µg/mL) of beauty products for different times (1, 4 h). This was done to evaluate how well TA protects against cytotoxicity induced by the examined products.

Our results revealed that TA extract significantly protects skin fibroblasts against the toxicity induced by all doses of beauty products at both times (1, 4 h) (Fig. 3a, b). However, the protective effect against toxicity induced by examined products was more pronounced when cells were preincubated with a safe dose (50 µg/mL) of TA extract for 48 h (Fig. 4a, b). Additionally, we extended the experiment by using preincubated skin fibroblasts Hs27 with a safe dose (50 µg/mL) of TA extract for 72 h, followed by exposure to beauty products for 1 and 4 h, to evaluate the protective effect against beauty product-mediated cytotoxicity. However, we did not find any increase in cell viability of skin fibroblasts after preincubation for 72 h. The reason might be the extortion of culture media for 72 h. Together, these results suggest that the TA extract can protect against the toxicity induced by beauty products, and it is most effective when skin fibroblasts are preincubated for 48 h with TA extract.

**Fig. 3.**
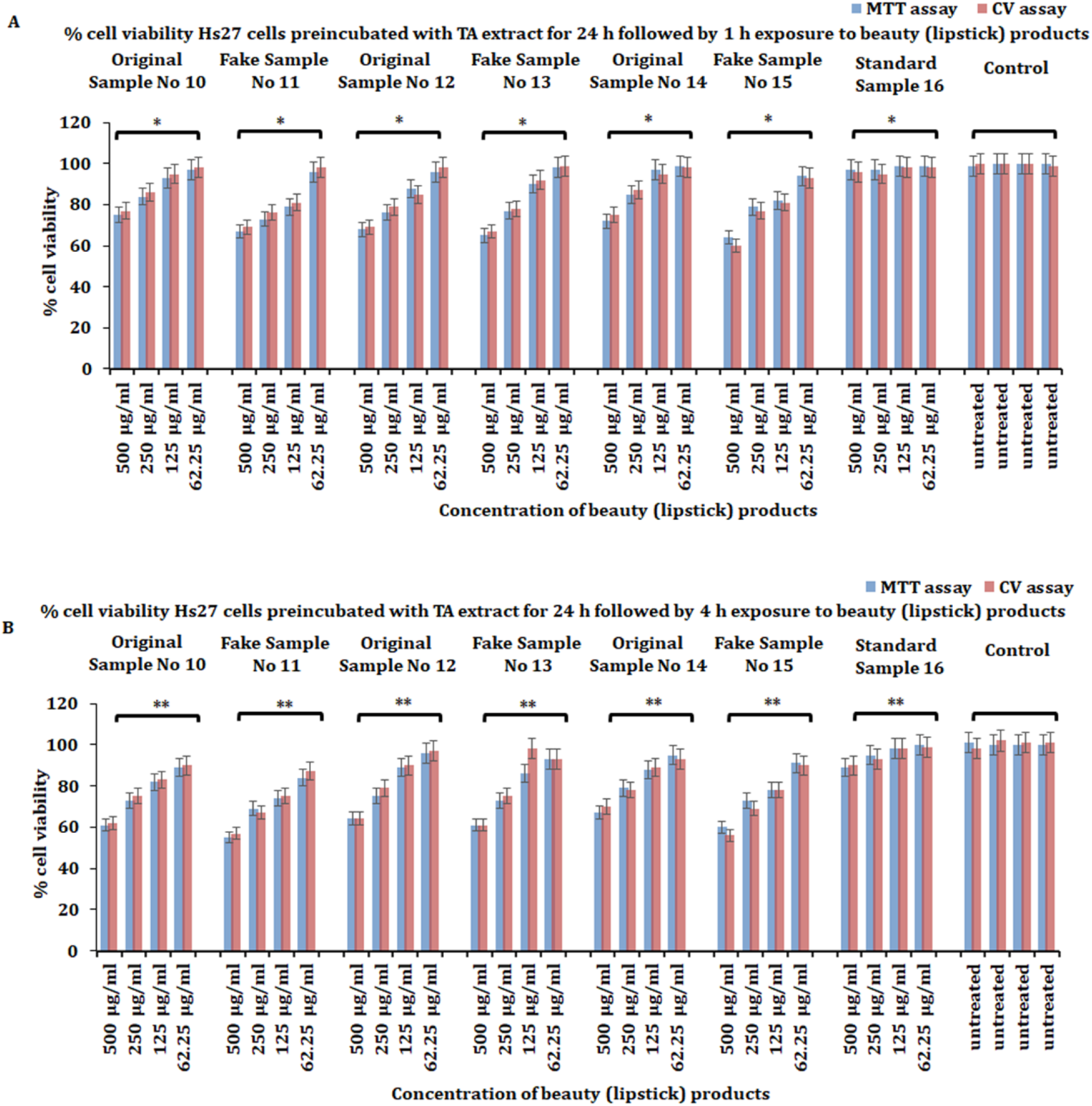
Protective effect of non-toxic dose (50 µg/mL) TA extract in Hs27 skin fibroblasts against the toxicity of personal care products. (**a**) Pre-treatment of Hs27 skin fibroblasts for 24 h with non-toxic dose (50 µg/mL) of TA extract, followed by treatment with varying doses (62.5, 125, 250, 500 µg/mL) of beauty (lipstick) products for 1 h to evaluate the percent cell viability. (**b**) Pre-treatment of Hs27 skin fibroblasts for 24 h with non-toxic dose (50 µg/mL) of TA extract, followed by treatment with varying doses (62.5, 125, 250, 500 µg/mL) of beauty (lipstick) products for 4 h to evaluate the percent cell viability. The data presented here is based on experiments done in triplicate and the mean value of ± SE. The p-value less or equal to 0.05 was considered to be statistically significant, \**p* ≤ 0.05, \*\**p* ≤ 0.01.

**Fig. 4.**
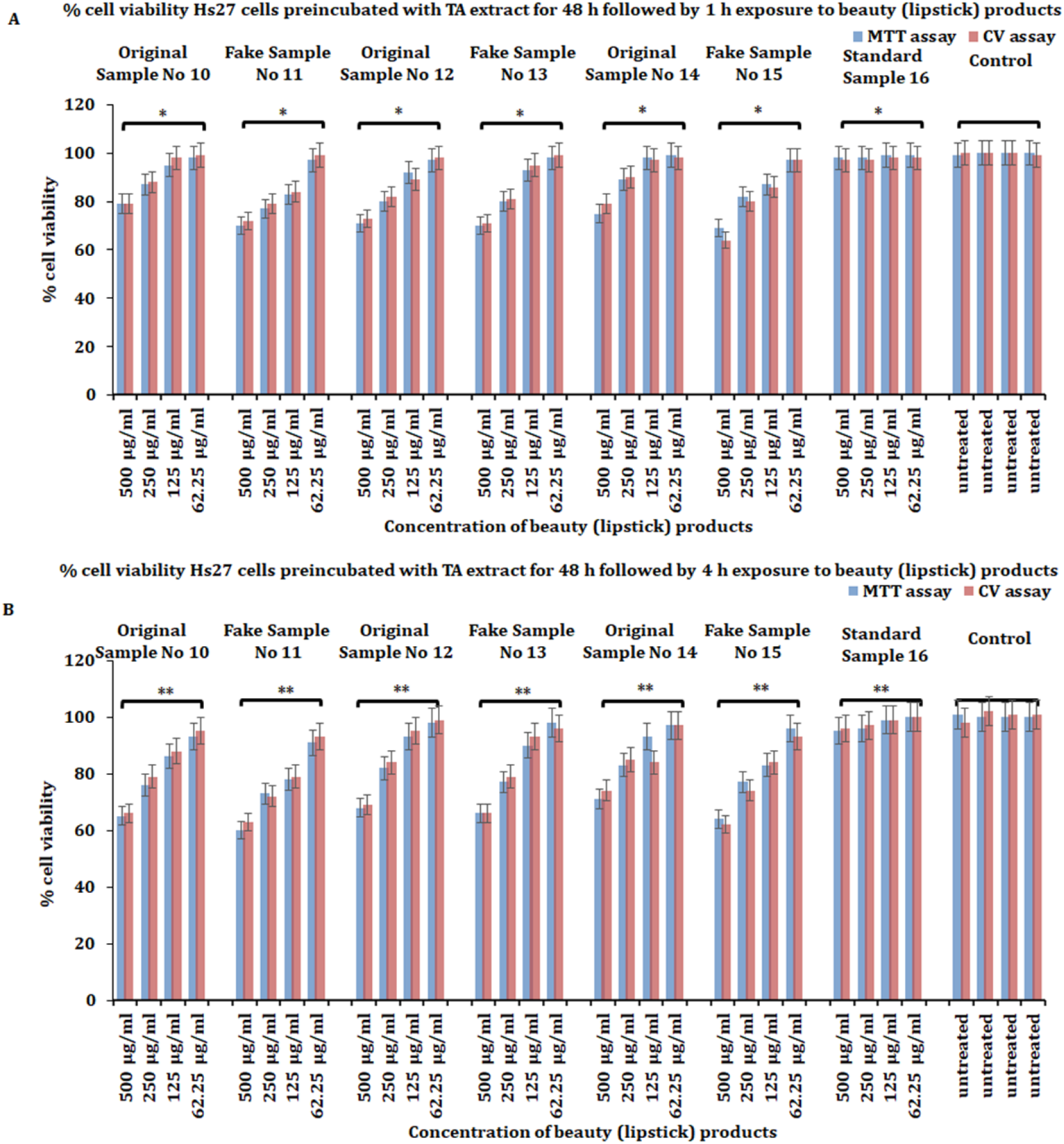
Protective effect of non-toxic dose (50 µg/mL) TA extract in Hs27 skin fibroblasts against the toxicity of personal care products. (**a**) Pre-treatment of Hs27 skin fibroblasts for 48 h with non-toxic dose (50 µg/mL) of TA extract, followed by treatment with varying doses (62.5, 125, 250, 500 µg/mL) of beauty (lipstick) products for 1 h to evaluate the percent cell viability. (**b**) Pre-treatment of Hs27 skin fibroblasts for 48 h with non-toxic dose (50 µg/mL) of TA extract, followed by treatment with varying doses (62.5, 125, 250, 500 µg/mL) of beauty (lipstick) products for 4 h to evaluate the percent cell viability. The data presented here is based on experiments done in triplicate and the mean value of ± SE. The p-value less or equal to 0.05 was considered to be statistically significant, \**p* ≤ 0.05, \*\**p* ≤ 0.01.

## Discussion

Exposure to numerous environmental pro-oxidants such as pollutants, radiation, chemicals, drugs, and cosmetic products causes oxidative stress in human beings, specifically at the cellular level in various organs including the skin and liver [29]. These pro-oxidant agents are harmful because they interact with bio-membrane proteins and lipids, which induces lipid peroxidation and interacts with DNA to induce genotoxicity in cells [21]. The cosmetic products market is one of the fastest-growing industries in the world and it is estimated that by 2022 it will earn more than $430 billion [22]. On average men and women use 6 and 12 cosmetic products, respectively, and every day in the United States [30]. More than 12,000 industrial and synthetic chemicals have been reported as used in the synthesis of cosmetic products. However, less than 20% of these chemicals are reported to have a safe toxicity profile and not much attention has been paid to the usage of cosmetic products derived from these synthetic chemicals [6]. Synthetic personal care products induce irritation, allergic rashes, and toxicity to skin cells by generating ROS through oxidative stress mechanisms. Excessive production of ROS by personal care products in skin fibroblasts causes a variety of skin diseases including melanoma and associated cancers [31].

The aim of the current study is to evaluate the toxicity profile against Hs27 skin fibroblasts of beauty products and their chemical analysis. Furthermore, we evaluated the protective effect of TA extract after Hs27 cells were exposed to varying doses of personal care products. Before the 1960s beauty products and personal care products had a good safety record except for a few make-up products which contained toxic chemicals and heavy metals such as cadmium, lead and mercury which were lethal to human beings [32]. Since the 1960s numerous scientific reports revealed that cosmetic products contain significant intoxicants in the form of chemicals and heavy metals which produce long-lasting photo-allergic and inflammatory skin reactions, subsequently posing a serious threat to consumers. For the past decade, the safety profile and chemical composition of synthetic personal care products have attracted great attention for evaluating the toxicity and photosensitization effects induced by these products against in-vitro skin fibroblast models and in vivo animal models.

Therefore, we set out to evaluate the chemical composition and cytotoxicity induced by beauty products against Hs27 skin fibroblasts. Our results revealed that a significant amount of cytotoxicity was induced by examined products when Hs27 skin fibroblasts were exposed to higher doses of the products for 1 and 4 h. To analyse the chemical analysis of the selected products, our results revealed that the cell-killing induced was due to the presence of a collective concentration of heavy metals in personal care products. Previous reports suggest that the toxicological data of some chemical compounds such as benzalkonium chloride and diazolidinyl urea induces cytotoxicity and activates ROS-mediated mitochondrial-dependent apoptosis in cells [33,34]. Avobenzone, commonly used in various personal care products promotes cell killing in human trophoblasts cells by calcium-mediated depolarization of the mitochondrial membrane potential to activate intrinsic apoptosis [35]. To evaluate whether the products in our study promote cytotoxicity by the activation of apoptosis, DNA fragmentation analysis demonstrated there was no ladder pattern of DNA fragments when Hs24 cell lysate samples exposed to different doses of personal care products were resolved by agarose gel electrophoresis. This strongly suggests that the cell-killing of Hs27 fibroblasts by personal care products was due to necrosis.

Chemical and heavy metal analysis is currently an important parameter for establishing the safety of cosmetic products [28]. Although heavy metals are present in trace amounts in cosmetic products, long-term usage and exposure to personal care products that contain heavy metals even in traces can accumulate and pose a danger to vital organs such as the liver, kidneys, etc [36]. Some heavy metals such as Cd, Ni, and Cr can act as carcinogens to induce different types of malignancies and are categorized as group 1 human carcinogens by the International Agency for Research on Cancer (IARC) [37]. Apart from causing malignancies, these heavy metals pose a greater risk of neurological and cardiovascular disorders [28]. Accumulation of As after entering through various routes into the body causes various problems such as malignancies in the gastrointestinal system, lungs, and urinary tract [38].

Intriguingly, our results revealed that no As was detected in personal care products investigated in the current study. Cd is another heavy metal found in personal care products and long-term exposure to these products causes serious damage to kidneys. The concentration of Cd in the samples ranged from 1-9 ppb which is well below the desired concentration [39]. Cr and its salts provide brightness to cosmetic products. Cr is very essential for regulating insulin function and helps in glucose metabolism [6]. In the current study we observed maximum concentration of Cr in branded samples as compared to fake samples but within the permissible concentration range. Pb exists in various cosmetic products and accumulates in the body either by inhalation or oral ingestion [40]. Accumulated Pb disturbs the central nervous system during fetus development in pregnancy and disturbs nerve transmission by interfering with calcium channels [40]. Pb detection concentration in the current study was also within the permissible range in all personal care products. Despite all the heavy metals in the current study found in permissible concentrations, due to the collective presence of all heavy metals in single personal care products and long-term exposure, they might pose a serious threat to human beings by causing various types of disorders. Owing to the presence of heavy metals, our cell viability assay suggests that these heavy metals collectively cause toxicity in Hs27 skin fibroblasts.

Products contain natural ingredients consisting of plant extracts, fruits, etc., that are rich in polyphenols and terpenes which act as antioxidants. The primary function of these compounds is to neutralize free radical species such as ROS, RNS, etc., and maintain the cell integrity free from any oxidative stress [41]. Mechanistically, these polyphenolic antioxidants possess phenolic groups that modulate protein phosphorylation of bio-membrane proteins to attenuate lipid peroxidation. They do this by acting as chain-breaking free radical scavengers thereby inhibiting oxidative reactions [42]. Plant extracts are in great demand in the cosmetics industry. The primary functions of plant products in the industry are to provide active ingredients and offer protection against other ingredients which can induce oxidative stress in skin fibroblasts [13,43].

Owing to the presence of abundant polyphenols and flavonoid compounds in TA extract, we intended to evaluate the protective effect of TA extract in Hs27 skin fibroblasts against the toxicity induced by personal care products. Our results demonstrated that preincubation with a non-toxic dose of TA for different periods of time (24, 48 h) followed by exposure to various doses (62.5, 125, 250, 500 µg/mL) of personal care products for 1 h and 4 h, indicated significant protection against personal care product-mediated cytotoxicity. The effect was more pronounced with 1 h exposure to personal care products when compared to 4 h. Together, these results suggest that the presence of abundant bioactive polyphenols with promising anti-scavenging activity in TA extracts significantly nullifies cytotoxicity promoted by personal care products in skin fibroblasts.

## Conclusion

In conclusion, this study has demonstrated that synthetic beauty products (lipstick products) exhibit significant cytotoxicity against in-vitro cellular model Hs27 fibroblasts. The toxicity to the in-vitro skin fibroblast model by synthetic products was due to the collective effect of traces of heavy metals present in beauty products. It is confirmed here that the cytotoxicity induced by the examined products was due to necrosis. Additionally, we observe TA extract, which is rich in antioxidant compounds offers a protective effect and neutralizes any toxicity of skin fibroblasts induced by beauty products. Owing to the presence of toxic chemicals and heavy metals that trigger various adverse effects on the skin and other vital organs, there is an urgent need to assess the safety of beauty products. This can be done by incorporating natural plant extracts into the formulations of personal care products to neutralize any adverse effects.

## Author Contributions

A. M. A, and F.M.A.Z designed and performed the experiments Visualization, data curation, acquisition, formal analysis, writing, and editing. A. M. A.: review and editing. B. R.: Conceptualization, design experiments, writing, editing, and making original draft of the manuscript.

## Institutional Review Board Statement

Not applicable

## Informed Consent Statement

Not applicable

## Data Availability Statement

Not applicable

## Acknowledgments

Researchers would like to thank the Deanship of Scientific Research, Qassim University for funding publication of this project.

